# Selection History Models in a Population under Ongoing Directional Selection

**DOI:** 10.1101/2025.11.11.687909

**Authors:** Anne C.M. Jansen, Mario P.L. Calus, Yvonne C.J. Wientjes

**Affiliations:** Animal Breeding and Genomics, Wageningen University and Research, 6700 AH, Wageningen, The Netherlands

## Abstract

The aim of animal breeding is to select the genetically best animals in the current generation to improve the performance of future generations for a specific breeding goal. With the continuous shift in breeding goals towards more balanced breeding, new traits may become of interest. Knowledge of the (indirect) selection history of these traits would be insightful before a trait is included in the breeding goal. Two models, BayesS and Ĝ, have been developed to assess the selection history of traits. BayesS estimates a parameter (*s*) that reflects the relationship between estimated additive effects and minor allele frequency, while Ĝ calculates the expected genetic change of a trait based on allele frequency changes and estimated additive marker effects. The aim of this study was to evaluate the performance of estimating *s*-values (based on BayesS) and Ĝ in an animal breeding context, focusing on their ability to detect selection for a trait with low heritability. Both Ĝ and *s*-value estimation were applied to a simulated dataset of a commercial pig breeding program under phenotypic selection, with varying heritabilities (0.05, 0.1, 0.3) and 30 generations of ongoing selection. Overall, both models were able to detect selection, where higher heritabilities and a larger sample size (for *s*-value estimation) or a larger selection interval (for Ĝ) resulted in increased detection of selection. The preferred model to identify selection varied based on the available data of the breeding population.

## Introduction

The aim of animal breeding is to select the genetically best animals in the current generation to improve the performance of future generations for a specific breeding goal. However, direct selection on breeding goal traits can cause indirect selection on traits not included in the breeding goal. The traits included in the breeding goal are determined based on societal and consumers’ demands. In the past, the focus of animal breeding programs was mainly on production traits, while currently there is more focus on balanced breeding, in which the aim is to simultaneously improve productivity, efficiency, environmental impact, animal health and welfare, food quality and safety, and genetic diversity (Olesen *et al*., 2000; Neeteson-van Nieuwenhoven *et al*., 2013; Boichard *et al*., 2015). With this change in breeding goal, new traits have become of interest. Knowledge of the (indirect) selection history of these traits would be insightful before the trait is included in the breeding goal. Strong ongoing indirect selection on a trait would be an indication that this trait has a non-zero genetic correlation to the other breeding goal traits. Understanding the history of selection on these traits is crucial to monitor unforeseen changes caused by indirect selection pressures.

Two models have been proposed that can assess the selection history of a trait using allele frequency data and estimated additive marker effects: BayesS and Ĝ (Beissinger *et al*., 2018; Zeng *et al*., 2018). BayesS estimates a parameter (*s*) that reflects the relationship between estimated additive marker effects and minor allele frequency (MAF) (Zeng *et al*., 2018). Those estimated additive marker effects reflect the average change in the phenotype when one allele at a locus is replaced by the other segregating allele. If a trait is under selection, the selection pressure will be strong on alleles with a large effect, thereby increasing the frequency of favourable alleles and decreasing the frequency of unfavourable alleles (Pritchard, 2001). Under positive selection, the frequency of loci with large effects is increased in the population and these are therefore more common. Therefore, a positive value for *s* is expected for this situation. If a trait is under negative selection, the allele frequency of large effect loci will be reduced in the population. This results in large effect loci with low MAFs, which is reflected by a negative value for *s* (Zeng *et al*., 2018; Ashraf and Lawson, 2021).

Ĝ uses the change in allele frequency between different generations of all loci combined with the estimated additive marker effects to evaluate the direction of selection (Beissinger *et al*., 2018). This method calculates the expected response to selection for all separate loci by multiplying the allele frequency change of a locus by its estimated additive effect. These values per locus are combined to get a genome-wide estimate for the response to selection of the past (Beissinger *et al*., 2018). It is expected that the allele frequency will change proportionally to the size and in the direction of the estimated additive marker effects (Wright, 1937; Beissinger *et al*., 2018).

BayesS was originally described to detect selection in human populations, which undergo strictly natural selection (Zeng *et al*., 2018). The model, or equivalent approaches, have occasionally been applied to artificially selected populations (i.e., tilapia, maize, cattle, and pigs) (Joshi *et al*., 2021; Palali Delen *et al*., 2023; Neshat *et al*., 2023; Zhang *et al*., 2025). However, detailed studies on how well this model works for finite populations under strong selection are still missing. Ĝ has been applied to breeding populations in different studies, resulting in the significant detection of selection in plants (Morales *et al*., 2021; Tilhou *et al*., 2022) and pigs (Chen *et al*., 2022). These studies investigated selection for highly heritable traits over various selection intervals. However, we want to determine whether there has been indirect selection on novel traits through selection on the breeding goal, for which the selection accuracy is lower and would be comparable to phenotypic selection for low heritability traits. Therefore, our interest was in the performance of the model for lower heritabilities.

The aim of this study is to evaluate the performance of estimating *s*-values (based on BayesS) and Ĝ in an animal breeding context, focusing on their ability to detect indirect selection for a trait or for a trait with a low heritability. Moreover, the aim is to identify the required amount of information that is needed to reliably determine the presence of selection. Both Ĝ and *s*-value estimation were applied to a simulated dataset of a commercial pig breeding program under phenotypic selection, with varying heritabilities and number of generations under selection to investigate their performance. The outcomes provide insights into the strengths and limitations of both these approaches and their potential for application in animal breeding programs to identify previous selection for traits, which may have been indirect by nature, based on estimated additive effects and allele frequency data.

## Material and Methods

### Simulated population structure

We simulated a commercial breeding population of pigs. First, a historical population was simulated to create genotypes for the base population using QMSim (v.2.0) (Sargolzaei and Schenkel, 2009) (Figure 1). The historical population started with 1,200 animals (600 males and 600 females). These animals were randomly selected and mated for 5,000 discrete generations. From the last generation of the historical population, 50 males and 500 females were randomly selected and randomly mated to create a pig line. To establish the line, random mating was continued for 100 generations while randomly selecting 50 males and 500 females each generation. The litter size was set to 6, consisting of 4 females and 2 males. The simulations were set up to generate an allele frequency pattern and linkage disequilibrium comparable to that in pig breeding populations.

**Figure 1.**
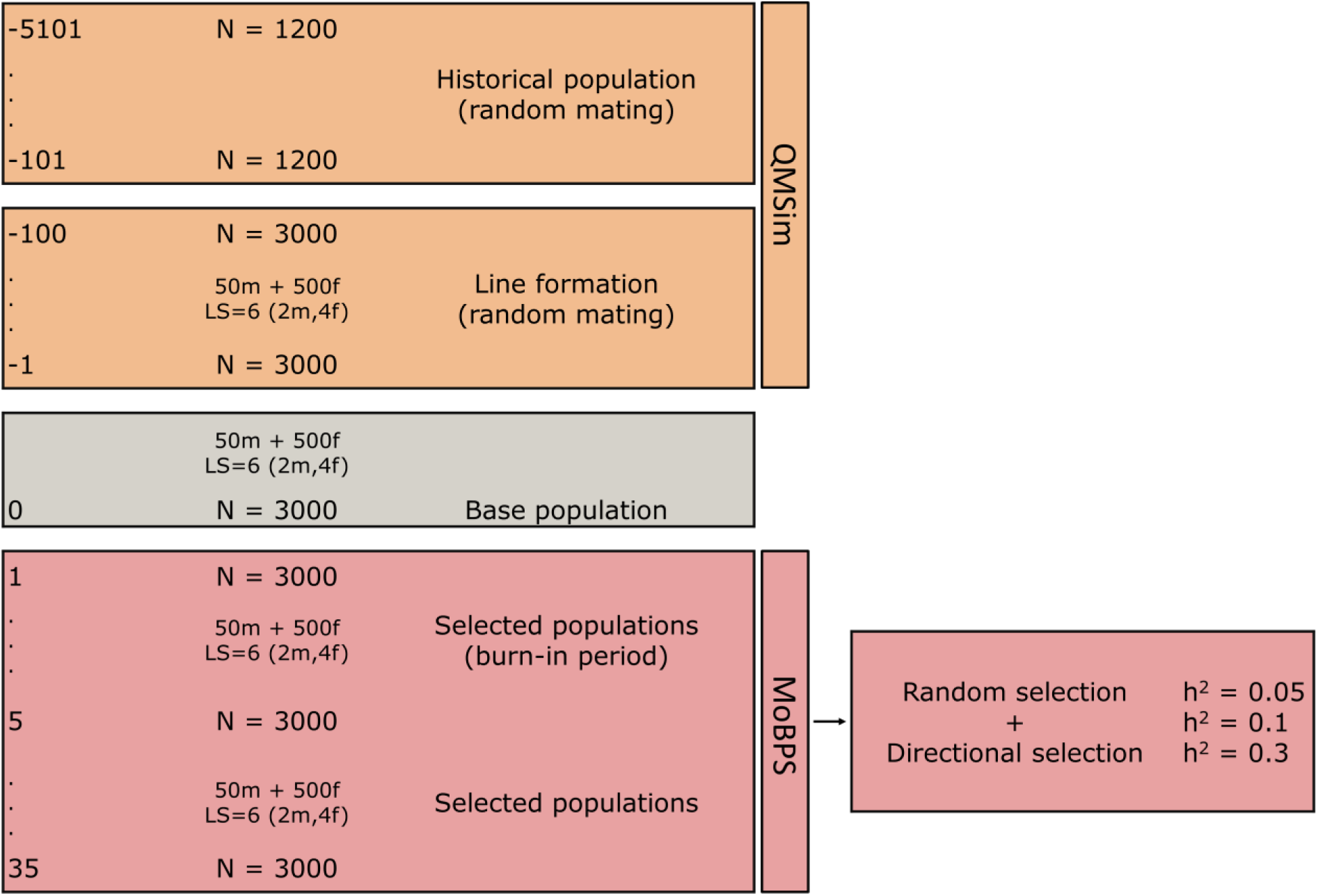
Overview of pig breeding simulation.

After line formation, the data was further processed using R (v.4.2.3) (R Core Team, 2023) and MoBPS (v.1.11.53) (Pook *et al*., 2020). From the last generation of line formation, 50 males and 500 males were randomly selected and mated to become parents of the base population. Thereafter, 2 different selection strategies were applied, namely directional selection based on own performance and random selection. A burn-in period of 5 generations was used to establish population structure, where the same selection strategy was applied as for that replicate, after which selection was continued for 30 generations more. In total, we used 50 replicates, and in each replicate a unique base population was created. To ensure comparability between selection scenarios, the same base population and seed were used for the different selection strategies within a replicate. This means that the same causal loci were sampled for the random and directional scenarios and for the three different heritability scenarios within each replicate. This was done to make sure that the observed differences between the random selection scenarios were strictly because of the difference in heritability and not due to sampling differences, while the differences between the directional selection scenarios were because of the differences in heritability and selection.

### Genome size

Just like the pig genome, 18 chromosomes were simulated, and the map length of each chromosome was set to be similar to that of pigs (Tortereau *et al*., 2012). Each chromosome was assigned 20,000 loci in the first generation of the historical population that were randomly spaced and had a randomly sampled allele frequency. The recurrent mutation rate was 2.5 × 10^−5^ in the historical population. From the last generation of the historical population, loci with a minor allele frequency (MAF) higher than 0.01 were selected and no new mutations were introduced after this point. After line formation, the allele frequency distribution of segregating loci in the base population was slightly U-shaped (Figure 2), which is caused by genetic drift and is in line with the expectations for random mating populations (Walsh and Lynch, 2018), and is also seen for whole-genome sequence data in livestock (Daetwyler *et al*., 2014; Brøndum *et al*., 2014; Heidaritabar *et al*., 2016). The linkage disequilibrium (LD) decay in the base population (Figure 3) was comparable to the LD in commercial pig breeding populations (Badke *et al*., 2012; Veroneze *et al*., 2013).

**Figure 2.**
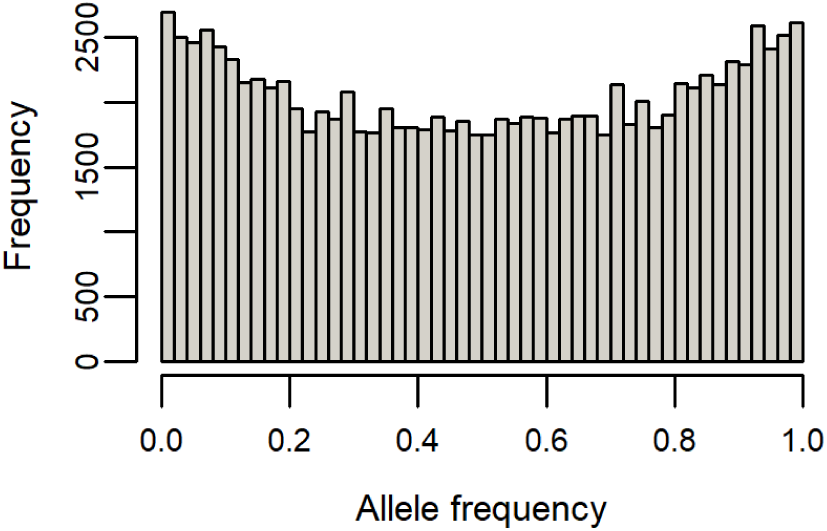
Allele frequency distribution in one replicate in the last generation of line formation.

**Figure 3.**
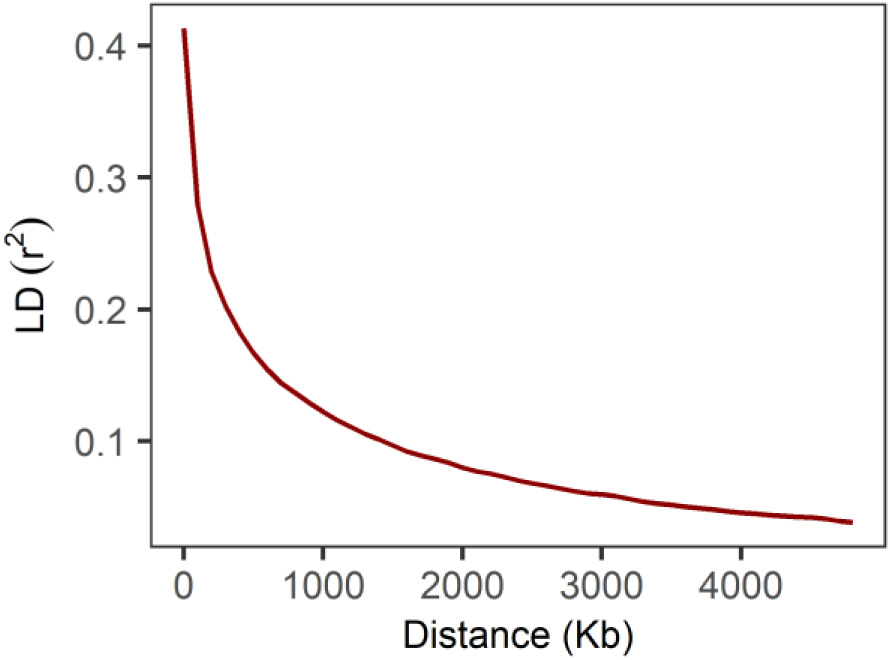
Linkage disequilibrium decay of one replicate in the last generation of line formation.

From the segregating loci in the base population, 2,000 loci were randomly sampled to become a causal locus. From the remaining segregating loci, 20,000 markers were selected by first dividing all loci into 100 bins based on their allele frequency, where bin 1 contained alleles with a frequency between 0 and 0.01, bin 2 between 0.01 and 0.02 and so on. From each bin, 200 loci were randomly sampled as markers, leading to a uniform allele frequency distribution for markers. This is representative of SNP arrays used in livestock (Matukumalli *et al*., 2009; Kranis *et al*., 2013).

### Phenotypes

Additive effects were sampled for all causal loci from a gamma distribution with shape parameter 0.4. The effects were randomly assigned a positive or negative sign, using the criteria that there were always 1,000 positive and 1,000 negative effects. The additive effects were multiplied by the allele counts of the causal loci (0, 1, or 2), to get the additive genetic values of each individual. The effects of the causal loci were scaled in MoBPS to ensure a mean of 0 and an additive genetic variance in the base population equal to the target heritability. Standardization of the additive genetic variance ensured that the causal effects were on a comparable scale across replicates. The target heritability (ℎ^2^) was set in the base population before the burn-in period, and was defined as 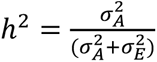, where 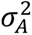 is the additive genetic variance and 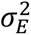 is the environmental variance. Based on the standardized additive genetic variance, the environmental variance was computed as 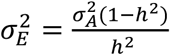, and was therefore different between different heritabilities. Environmental deviations were sampled from a normal distribution with mean 0 and variance 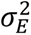. Phenotypes were then generated by adding the environmental deviations to the additive genetic values.

### Selection

Two forms of selection, i.e., random selection and directional selection were simulated for 30 generations after the 5-generation burn-in period. Moreover, three heritabilities (0.05, 0.1, and 0.3) were simulated. This resulted in 2x3 = 6 scenarios, with 50 replicates per scenario (Table 1). As the aim was to study the selection history of traits that have possibly been under indirect selection, we decided to apply phenotypic selection in the simulated dataset. The reason for this is that if selection has acted indirectly on traits, the accuracy would be lower than if, for instance, genomic selection had acted on these traits. If the models perform well on traits under phenotypic selection on a trait with a low heritability, it can be assumed that they could be applied to traits under indirect genomic selection as well.

**Table 1.**
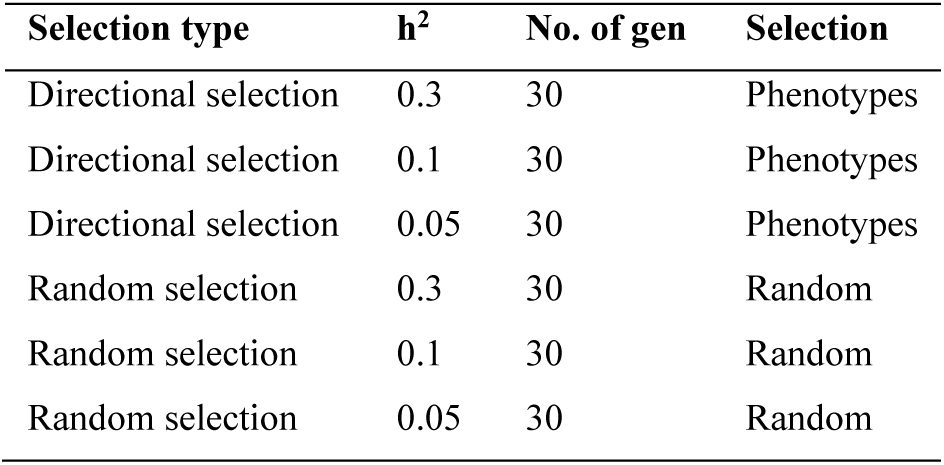
Overview of the selection strategies.

### Ĝ model for selection

The first model that we tested was the Ĝ model, developed to detect the selection history of traits based on the expected genetic gain, resulting from the change in allele frequency of markers and their estimated additive effect (Beissinger *et al*., 2018; Mahmoud *et al*., 2023). Ĝ is calculated as 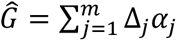, where Δ_𝑗_ is the change in allele frequency at locus *j*, 𝛼_𝑗_ is the estimated effect of this locus *j*, and *m* is the number of genotyped markers. The model uses the change in allele frequency between two timepoints or generations. In our simulation, we applied it to varying numbers of generations to detect selection over 1, 5, 10, 15, and 20 generations, looking back from generation 30 (i.e., to detect selection over 20 generations, the change in allele frequency was determined between generation 10 and 30). The SNP effects of segregating loci (i.e., 𝛼) of the SNP array were estimated in generation 30 using rrBLUP in MoBPS.

To assess if Ĝ is significantly different from zero, the value is compared to a permuted null-distribution, for which 1,000 permutations were used (Beissinger *et al*., 2018). The Ghat R-package was used to determine the number of independent markers based on LD in the starting generation, with a maximum window of 500kb and a maximum r^2^ of 0.03, as is recommended for this data structure (Mahmoud *et al*., 2023). This value was used to adjust the normal null distribution that resulted from the permutations as: 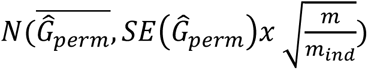, where *m* is the number of markers, and *m*_ind_ is the number of independent markers. Thereafter, the observed Ĝ is compared to this null distribution to get a *p*-value. The resulting *p*-value was considered significant when <0.05. A Ĝ value that is significantly greater than the permuted mean indicates positive directional selection, and a value significantly lower than the permuted mean indicates negative directional selection. When Ĝ was not significantly different from the permuted mean, there was no significant evidence for selection.

### *s-*value estimation

The second model aims to estimate a parameter *s* that reflects the relationship between the estimated additive effects of loci and their minor allele frequencies (MAF) at one timepoint in a population (Zeng *et al*., 2018). This relationship provides information about ongoing selection in a population. In case the MAF and additive effects are unrelated, the *s-*value would be zero. A value for parameter *s* < 0 reflects that the effects of rare loci are, on average, larger than the effects of common loci, which is seen when large effect loci are kept at low frequencies by negative selection. Contrarily, when *s* > 0, the common loci have larger effects than rare loci and this is expected in human populations when selection increases the frequency of large-effect loci by positive selection (Zeng *et al*., 2018; Ashraf and Lawson, 2021). However, artificial selection in finite populations results in a rapid increase of beneficial alleles. The beneficial alleles with large effects rapidly increase towards fixation and therefore only remain at intermediate frequencies for a short time. This results in loci with a large effect having a relatively low MAF. Alleles with smaller effects experience a lower selection pressure and remain at higher MAF. This results in a negative relationship between the MAF and effect size. Therefore, a negative value for *s* is expected for livestock populations under positive selection as well, whereas an *s*-value of zero would indicate that no selection has occurred.

A genomic restricted maximum likelihood (GREML) model was used to determine the value of *s* that best fitted the data. The following mixed model was used for the GREML analysis:

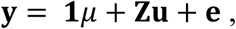

where **y** is a vector of phenotypes, 𝟏 is a vector of ones, 𝜇 is the overall mean, **Z** is a design matrix linking the phenotypes to the breeding values of the animals, 𝐮 is a vector of breeding values with 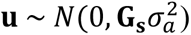 where **G_s_** is the genomic relationship matrix constructed using parameter *s*, and 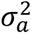 the total additive genetic variance. The residuals are represented by **e**, with 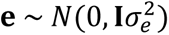, where **I** is the identity matrix and 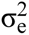 is the residual variance. The **G**-matrix with the parameter *s* is calculated as:

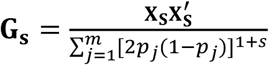

Where **X_s_** is a scaled genotype matrix, *m* is the number of loci, 𝑝_𝑗_ is the allele frequency of locus *j,* and parameter *s* ∈ [−1,1].

The element for individual *i* and locus *j* in the scaled genotype matrix **X_s_** is defined as:

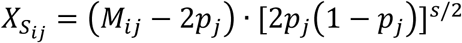

Where **M** is the allele count matrix with dimensions *n* x *m*, where *n* is the number of individuals and *m* is the number of loci, and 𝑀_𝑖𝑗_ is the allele count (0, 1, 2) for individual *i* at locus *j*. The notation is based on (Speed *et al*., 2017; Zeng *et al*., 2018).

We varied the values for *s* from -1 to 1 with increments of 0.1, resulting in 21 **G_s_-**matrices that were differently scaled. It is good to note that with *s* = 0, the resulting **G_s_** matrix was equivalent to the **G** matrix obtained with method 1 described in (VanRaden, 2008), and *s* = -1 resulted in the **G** matrix obtained with method 2 described in (VanRaden, 2008). The **G_s_** matrices were calculated using calc_grm (Calus and Vandenplas, 2016). The different **G_s_** matrices were then used in a GREML analysis using MTG2 (Lee and van der Werf, 2016), resulting in a log likelihood value for each considered *s-*value (Speed *et al*., 2017; Bouwman *et al*., 2017). The *s-*value that resulted in the highest log likelihood was considered to best fit the data. Using AIC scores, it was determined whether the model fit for those identified *s-*values substantially differed from the fit for the other *s-*values. The AIC score was calculated as 𝐴𝐼𝐶 = −2 𝑙𝑛(𝐿) + 2𝑘, where 𝑘 is the number of estimated parameters and 𝐿 is the likelihood estimate (Burnham and Anderson, 2003). Then we estimated for each tested *s-*value Δ𝐴𝐼𝐶_𝑠_ = 𝐴𝐼𝐶_𝑠_ − 𝐴𝐼𝐶_𝑚𝑖𝑛_, where 𝐴𝐼𝐶_𝑚𝑖𝑛_ is the model with the lowest AIC value (which is the best fitting model) and 𝐴𝐼𝐶_𝑠_ are the 𝐴𝐼𝐶 scores for each value for *s*. These values were evaluated based on the criteria from Burnham and Anderson (2003), where a Δ𝐴𝐼𝐶_𝑠_of 0-2 reflects that 𝐴𝐼𝐶_𝑚𝑖𝑛_ is not a substantially better fit than 𝐴𝐼𝐶_𝑠_, 4-7 reflects that 𝐴𝐼𝐶_𝑚𝑖𝑛_ is a considerably better fit than 𝐴𝐼𝐶_𝑠_, and >10 means that there is essentially no support for the 𝐴𝐼𝐶_𝑠_ value. Here, we therefore considered that Δ𝐴𝐼𝐶_𝑠_scores of >10 indicate that there is a substantially better fit for 𝐴𝐼𝐶_𝑚𝑖𝑛_. We identified across replicates how often the optimal *s*-value differed from zero with a Δ𝐴𝐼𝐶_𝑠_ score >10 and used this as the indication of selection.

### Development of selection signals

The development of the selection signal over the selected generations was further investigated to get insight into the number of generations of selection that are required before selection can be detected. Therefore, *s-*values were estimated in generations 1, 5, 10, 15, 25, and 30 (after the burn-in period of 5 generations). For each generation, the Δ𝐴𝐼𝐶_𝑠_ were calculated to assess the certainty of the *s-*value estimates, providing insight into how clearly the selection signal could be identified over time.

### Ascertainment bias of SNP sampling

In the simulations of the SNP arrays, loci were sampled in allele frequency bins, resulting in a uniform allele frequency distribution on the array. In contrast, the causal loci were randomly sampled from all segregating loci, which had a U-shaped allele frequency distribution. The sampling of the SNP array could therefore have resulted in ascertainment bias, where the SNP array is not a good representation of the allele frequency distribution of the causal loci. To investigate the impact of this on the estimation of the *s*-values, simulations with a heritability of 0.3 were repeated for both random and directional selection, where the SNPs for the array were selected completely at random, retaining the U-shaped allele frequency distribution. The *s-*values for these simulations were compared to the *s-*values for the simulations with a uniform SNP array. Furthermore, the *s-*values based on the 2,000 actual causal loci were determined and included in the comparisons.

### Development of MAF of causal loci by selection

To evaluate how the genetic architecture evolved under selection, we evaluated the correlation between MAF and effect size of causal loci in the last generation of selection. This correlation was expected to arise because of selection, as before selection this was zero. The regression coefficients of the absolute causal loci effect on the MAF were calculated in generation 30. The mean slope was calculated over the 50 replicates for heritabilities of 0.05, 0.1 and 0.3 for directional selection.

## Results

### Evidence of selection

To investigate the capability of the models to identify selection in a pig breeding program, 30 generations of selection were simulated, preceded by 5 generations of selection as a burn-in period (Figure 4). The average breeding values after a total of 35 generations of selection for directional selection were on average 2.25, 4.04, and 10.03 for heritabilities of 0.05, 0.1, and 0.3, respectively. For random selection, the breeding values in the last generation were on average -0.07, -0.10, and -0.17 for heritabilities of 0.05, 0.1, and 0.3, respectively. This shows that for directional selection, the breeding value increased more with increasing heritabilities. For random selection, there has been no development of the breeding values, as expected.

**Figure 4.**
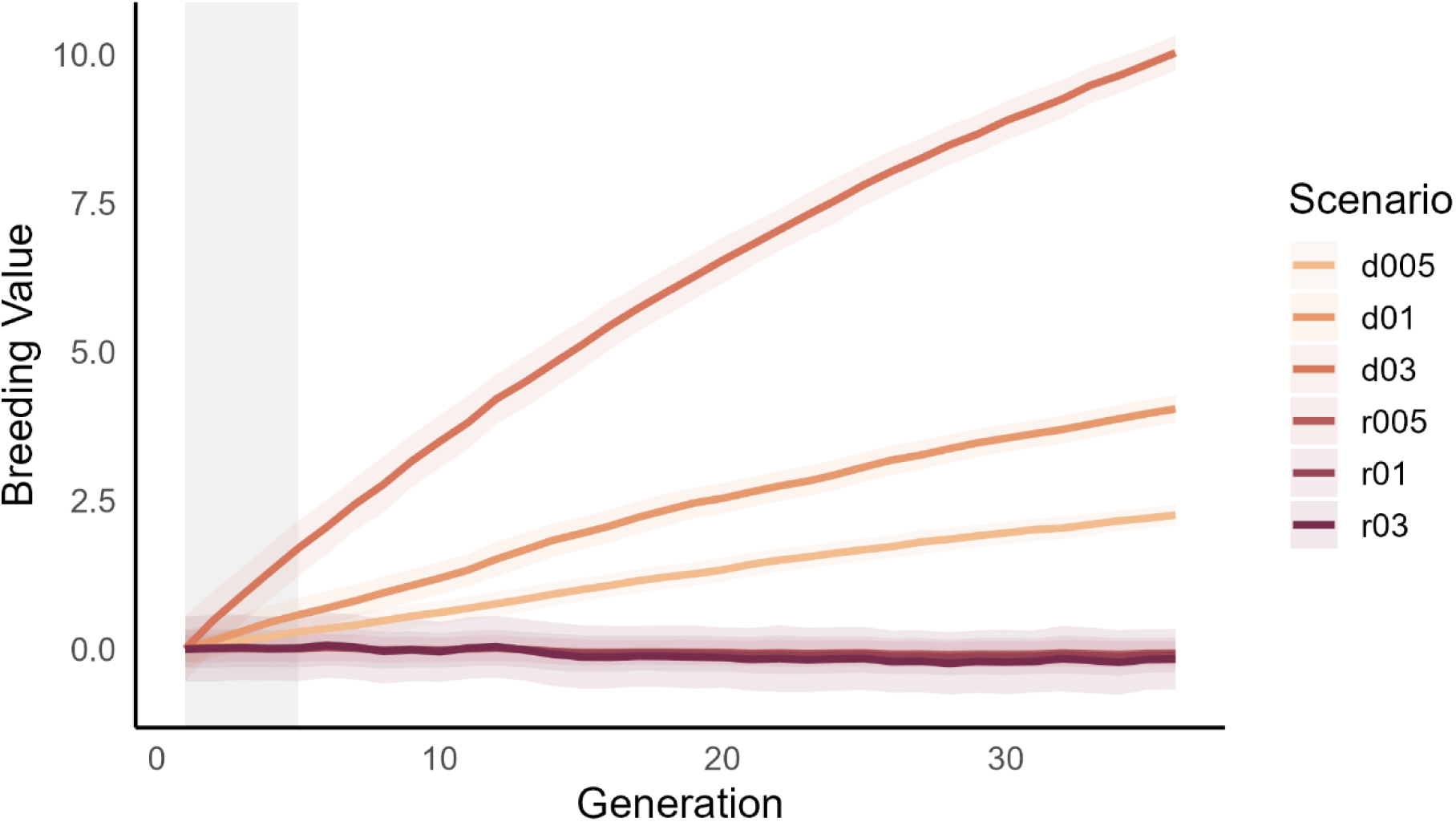
Trait development over generations for all selection scenarios: directional (d) and random (r) selection for heritabilities 0.05, 0.1, and 0.3. The burn-in period of 5 generations is shaded in grey.

### Ĝ model for selection

The first model that was tested to detect selection history was Ĝ, which was applied to five selection intervals, i.e., the number of generations between the two generations used for calculating the change in allele frequency (20, 15, 10, 5, 1 generations), while generation 30 was always the last generation. Ĝ never detected a significant signal for selection for the random selection scenarios, as was expected (Table 2). For the trait under directional selection, Ĝ was able to detect a significant signal of selection in many cases, and a higher heritability resulted in a higher probability of significantly detecting selection (Figure 5). Increasing the selection interval resulted overall in correctly identifying the selection more often. For a selection interval of 10 (i.e., gen. 20-30), selection was detected correctly for 98% of the replicates for h^2^=0.3, for h^2^=0.1 this was 56%, and for h^2^=0.05 this was 30%. However, increasing the selection interval further (i.e., for gen. 15-30 and 10-30) did not result in a strong increase in the indication of selection.

**Figure 5.**
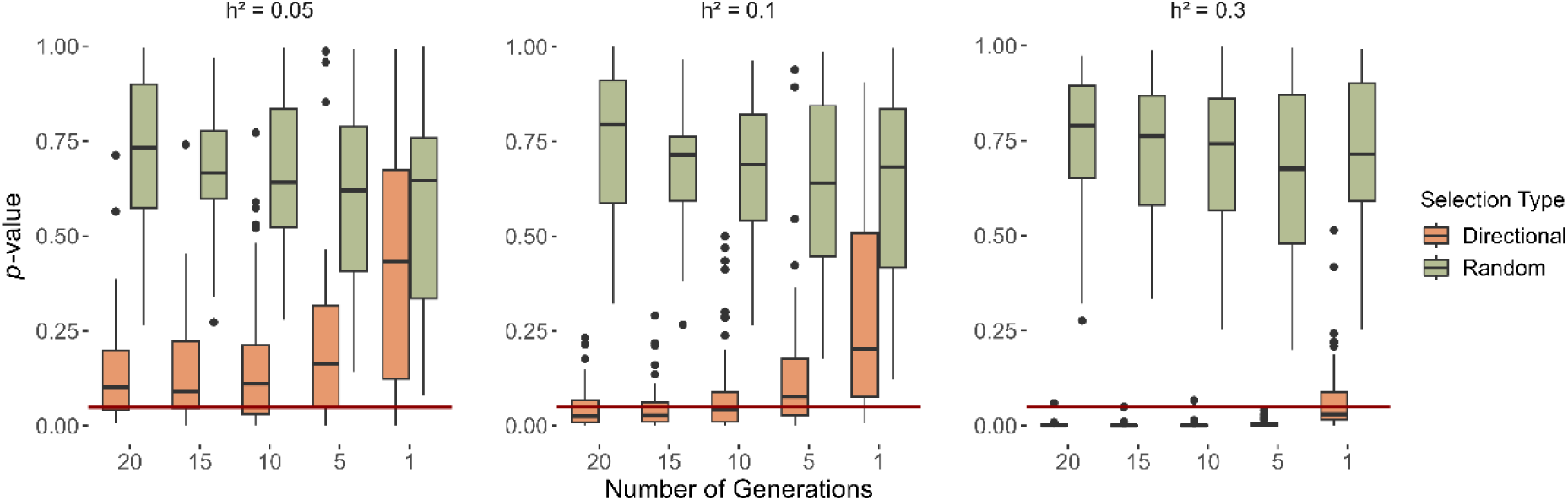
Results for Ĝ obtained for different selection types (directional and random selection), different heritabilities (0.05, 0.1, 0.3) and different number of generations of selection (20, 15, 10, 5, 1).

**Table 2.** Overview of number of replicates (out of 50) with a *p*-value < 0.05 for Ĝ.

|  | Directional selection |  |  | Random selection |  |  |
| --- | --- | --- | --- | --- | --- | --- |
| | $h^2 = 0.05$ | $h^2 = 0.1$ | $h^2 = 0.3$ | $h^2 = 0.05$ | $h^2 = 0.1$ | $h^2 = 0.3$ |
| gen. 10-30 | 16 | 34 | 49 | 0 | 0 | 0 |
| gen. 15-30 | 14 | 35 | 50 | 0 | 0 | 0 |
| gen. 20-30 | 15 | 28 | 49 | 0 | 0 | 0 |
| gen. 25-30 | 13 | 19 | 50 | 0 | 0 | 0 |
| gen. 29-30 | 5 | 10 | 31 | 0 | 0 | 0 |

### *s-*value estimation

The second model estimated *s-*values as an indication of selection, which were estimated for datasets containing generations 30, 29-30, or 25-30. For the scenarios with directional selection, the optimal *s-*value was on average between -0.8 and -1 for different heritabilities (Figure 6). Overall, an *s-*value of -1 had the highest likelihood with directional selection, except for the scenario with a heritability of 0.05 and using data of generation 30 only, where the optimal *s-*value was -0.8 on average.

**Figure 6.**
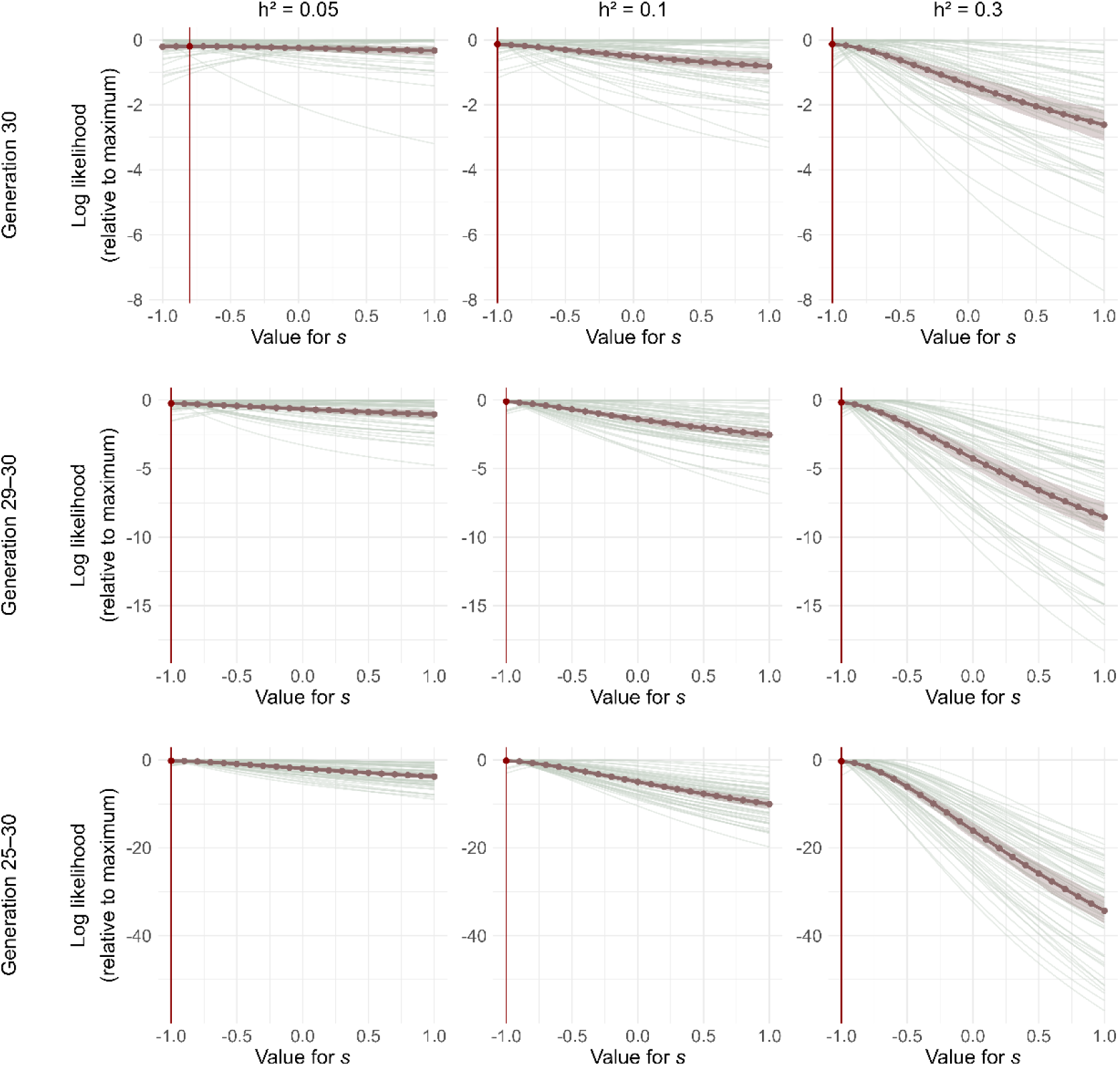
Log likelihood values for *s-*values ranging from -1 to 1 for directional selection. The thin lines represent the 50 replicates, where the thick line is the average of these replicates. The *s-*value that has, on average, the best likelihood value is highlighted in red.

The Δ𝐴𝐼𝐶_𝑠_scores revealed that for the *s-*values that were determined based on data from generation 30 only, there were no replicates that reached Δ𝐴𝐼𝐶_𝑠_ scores >10 compared to an *s*-value of zero, for any of the tested heritabilities (Appendix I, Table 3). This shows that the difference between the likelihood values for *s-*values that were determined to be best was not substantially better than *s-*values of zero. Enlarging the dataset by including the last two generations in the *s*-value estimation for a heritability of 0.3, resulted in 19 out of 50 Δ𝐴𝐼𝐶_𝑠_ scores >10 compared to an *s*-value of zero. Further enlarging the dataset resulted in Δ𝐴𝐼𝐶_𝑠_scores >10 for all heritabilities, and Δ𝐴𝐼𝐶_𝑠_scores >10 was reached for 2, 23, and 50 replicates for a heritability of 0.05, 0.1, and 0.3, respectively. Generally speaking, with increasing heritabilities and increasing number of generations of data, the differences between log likelihood values of the best fitting *s-*value and the other *s-*values increased.

**Table 3.** Overview of the number of replicates (out of 50) with a ΔAIC_s_ >10 for an *s*-value of zero.

|  | Directional selection |  |  | Random selection |  |  |
| --- | --- | --- | --- | --- | --- | --- |
| | $h^2 = 0.05$ | $h^2 = 0.1$ | $h^2 = 0.3$ | $h^2 = 0.05$ | $h^2 = 0.1$ | $h^2 = 0.3$ |
| gen. 30 | 0 | 0 | 0 | 0 | 0 | 0 |
| gen. 29-30 | 0 | 0 | 19 | 0 | 0 | 0 |
| gen. 25-30 | 2 | 23 | 50 | 0 | 1 | 1 |

For random selection, the optimal *s-*value was on average between -0.2 and -0.3 (Figure 7). The difference in the log likelihood values between the optimal *s-*value and the other *s-*values increased for higher heritabilities and an increasing number of generations of data provided, as was also seen under directional selection. The Δ𝐴𝐼𝐶_𝑠_scores revealed there were some replicates for the analysis of generation 30 only, where the best fitting *s-*value reached values >10 compared to other *s*-values (Appendix II). Looking at two and five generations of data, the number of replicates that reached an Δ𝐴𝐼𝐶_𝑠_score >10 compared to other *s-*values increased. However, when comparing the optimal *s-*value to an *s*-value of zero, Δ𝐴𝐼𝐶_𝑠_>10 was only reached twice: once for a heritability of 0.1 and once for 0.3, with the use of the last five generations of data (Table 3).

**Figure 7.**
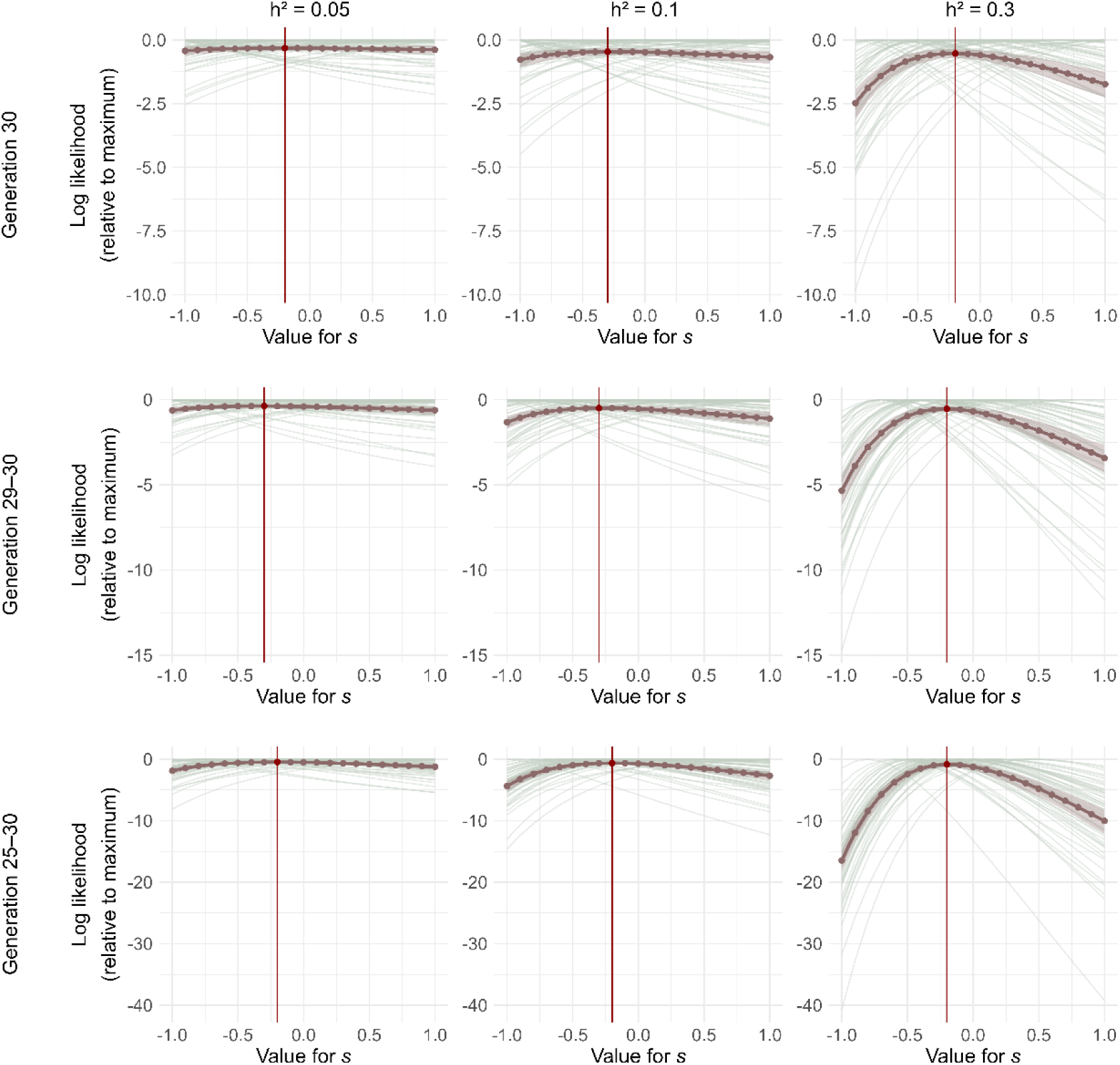
Log likelihood values for *s-*values ranging from -1 to 1 for random selection. The thin lines represent the 50 replicates, where the thick line is the average of these replicates. The *s-*value that has, on average, the best likelihood value is highlighted with a red vertical line.

### Development of selection signals

*s-*values were estimated in generation 1, 5, 10, 15, 25, and 30 for directional selection to investigate the development of the signal to detect selection. For heritability 0.3, the estimated *s-*values became more negative for an increasing number of generations of selection. In generation 1 (the first generation after the 5 generation burn-in period), the optimal *s-*value was on average determined to be -0.6, whereas in generation 10 and onwards this was estimated to be -1 (Figure 8). The Δ𝐴𝐼𝐶_𝑠_ scores reached values >10 with at least one other *s*-value in any generation, where in most cases this significance was reached when the optimal *s-*value was (close to) -1. For this heritability of 0.3, the highest Δ𝐴𝐼𝐶_𝑠_ scores and therefore the strongest selection signal, was found between generations 10-20. After this generation, the signal became weaker. Similar to heritability 0.3, the (on average) optimal *s-*value became more negative over time for a heritability of 0.1. Here, the optimal *s-*value was estimated at -0.4 in generation 1 and reached -1 in generation 30. For heritability 0.05, the optimal *s-*values did not develop uniformly in one direction as was seen for the other heritability values, and overall, the optimal *s-*values ranged between -0.1 and -1. For heritabilities of 0.05 and 0.1 the Δ𝐴𝐼𝐶_𝑠_scores only rarely reached values >10 with at least one other *s*-value and therefore the differences in model fit between the *s-*values were not substantial (Appendix III).

**Figure 8.**
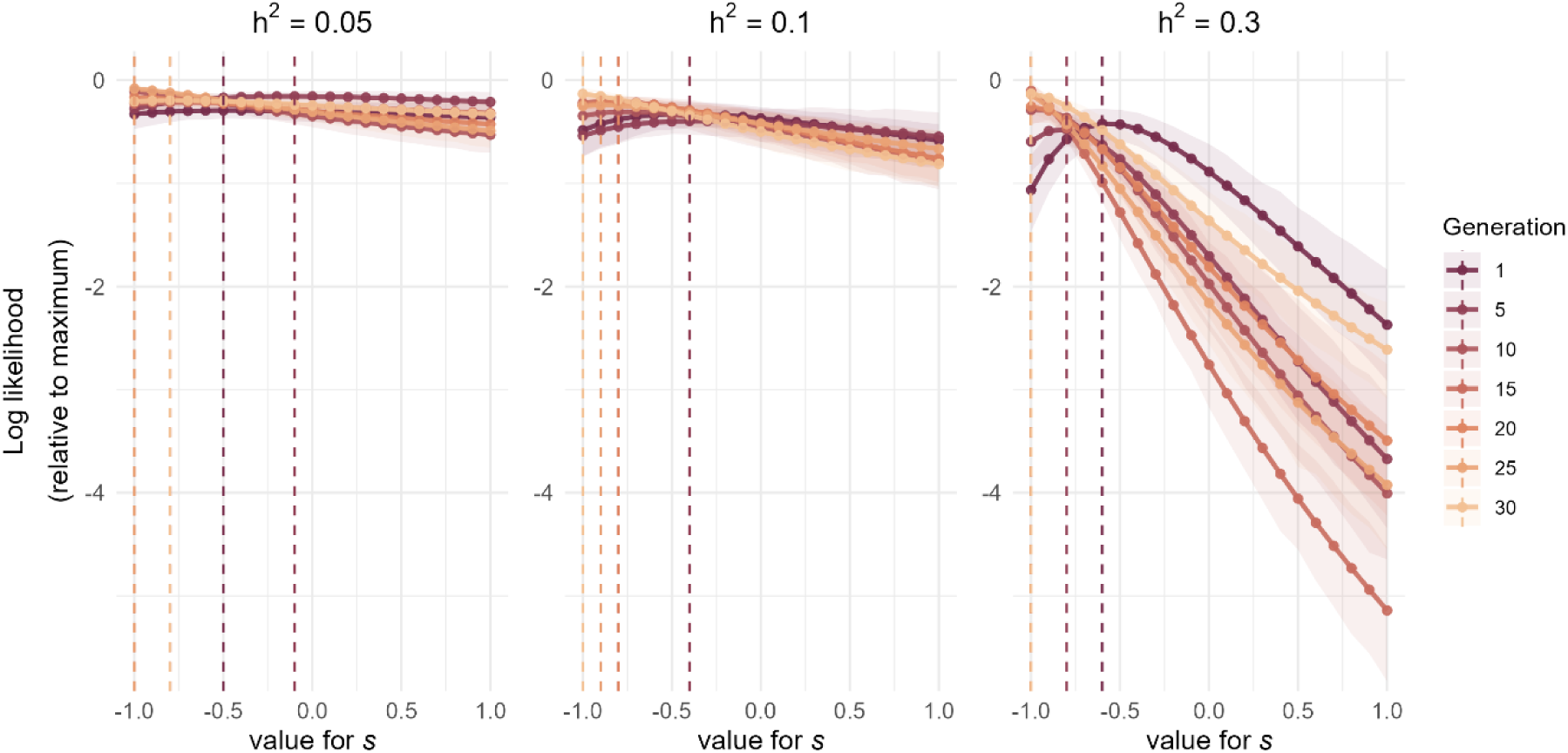
Log likelihood values for *s-*values ranging from -1 to 1 for directional selection, for generations 1, 5, 10, 15, 20, 25, and 30. The lines represent the average log-likelihood values for each population. The *s-*value that has, on average, the best likelihood value is highlighted with a vertical line.

### Ascertainment bias of SNP sampling

To determine if the design of the SNP array had an influence on the estimated *s-*values, the optimal *s-*values were also determined for a SNP array with U-shaped allele frequency distribution and compared to the original array with a uniform allele frequency distribution. Moreover, the *s-*values were determined directly for the causal loci. Under directional selection, the optimal *s-*value for the U-shaped as well as the uniform allele frequency SNP arrays was estimated to be -1 (Figure 9). Yet, for the causal loci the optimal *s-*value was estimated at -0.7. For random selection, the *s-*values for the U-shaped and uniform SNP arrays were estimated at -0.3 and -0.2, respectively. Where for the causal loci in this case, it was estimated to be 0.1.

**Figure 9.**
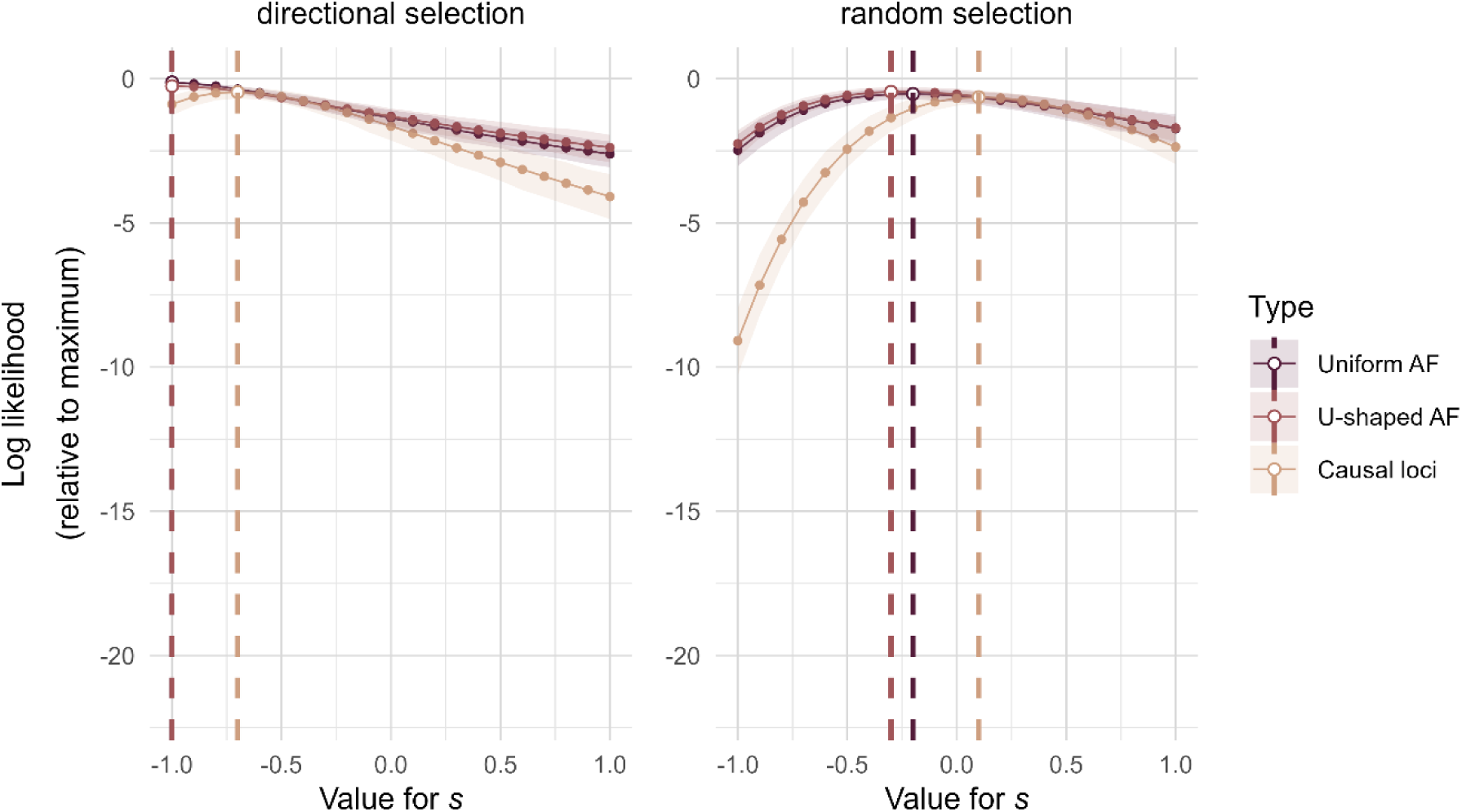
Log likelihood values for *s-*values ranging from -1 to 1 for both directional and random selection, with a heritability of 0.3, after 30 generations of selection. The lines represent the average log-likelihood values for a population with a different array; u-shaped allele frequency SNP array, uniform allele frequency SNP array, only the causal loci. The *s-*value that gave on average the optimal likelihood value is highlighted with a vertical line.

### Development of MAF of causal loci by selection

To evaluate the actual relationship between MAF of the causal loci and their effect sizes under selection, we examined the regression coefficients for the MAF of causal loci with their effect sizes in generation 30 of directional selection. The mean regression coefficients were calculated for heritabilities of 0.05, 0.1, and 0.3 to be -0.297, -0.430, and -0.512. So, the slope became more negative respectively when the heritability was higher (Figure 10). As for the random selection scenarios, for which the same causal loci were sampled, there is one common slope for the three heritabilities, where the mean slope is 0.0244.

**Figure 10.**
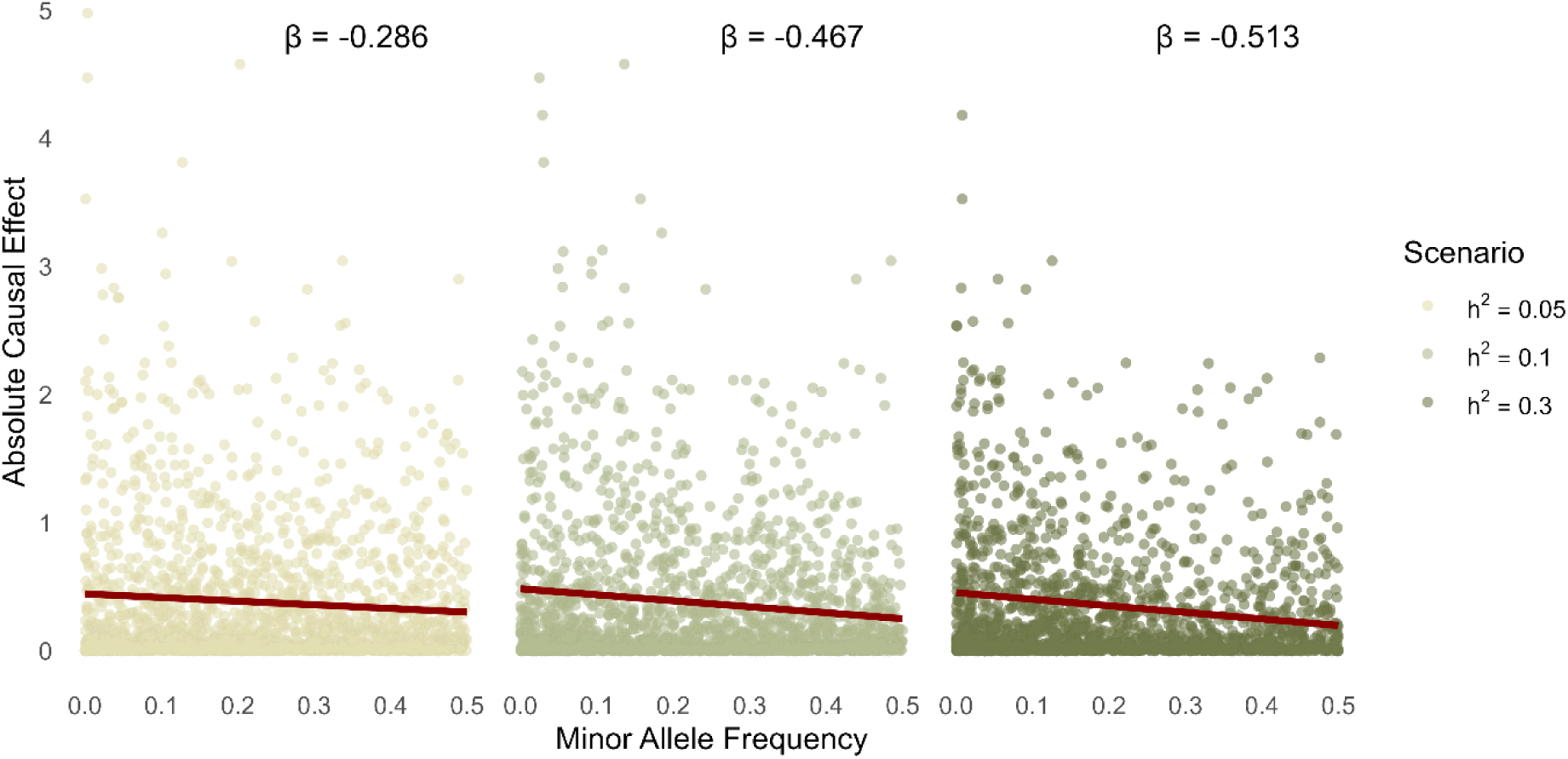
The effect of the causal loci versus their minor allele frequency with the regression slope for directional selection for all three heritabilities for 1 replicate in generation 30.

## Discussion

The aim of this study was to evaluate the performance of Ĝ and estimated *s-*values to investigate the selection history of traits in livestock populations. The models were applied to a simulated pig breeding program with either random or directional phenotypic selection for traits with heritabilities of 0.05, 0.1, and 0.3. Results showed that Ĝ was able to detect selection very accurately for a heritability of 0.3 and a selection interval of at least five generations. The accuracy of detection decreased for heritabilities of 0.1 and 0.05. Yet, Ĝ never detected a significant signal for random selection. For the estimation of *s-*values, a value of zero reflected no selection, and a deviation from zero with Δ𝐴𝐼𝐶_𝑠_ >10 was considered a significant indication of selection. Under directional selection, optimal *s-*values were generally estimated close to -1, with higher ΔAIC scores and more frequent strong deviations from zero for higher heritabilities and when more generations of data were included. For random selection, optimal *s*-values were slightly below zero, and only in some rare cases the optimal *s*-value was a considerable better fit to the data than an *s*-value of zero.

### Relating the simulations to real breeding populations

In our simulation, we applied 30 generations of selection. This number was chosen to represent the long-term selection that has been applied in commercial pig breeding programs. The nucleus populations of breeding companies have undergone continuous selection for meat production and reproductive traits for several decades, corresponding to around 30 discrete generations of selection (Amills *et al*., 2010; Merks *et al*., 2012). The selection history simulated here is therefore broadly comparable to that of real pig breeding populations.

This study has focused on the detection of selection in traits with low to moderate heritabilities (0.05, 0.1, and 0.3). Traits with heritabilities in the range of 0.05 to 0.1 in pigs represent common reproductive traits (i.e., total number born, number born alive, number born dead, number of stillborn piglets, number of mummies, litter weight, and piglet birth weight) (Schneider *et al*., 2012; Ye *et al*., 2018; Sell-Kubiak, 2021; Sell-Kubiak *et al*., 2022; Ogawa *et al*., 2022), while traits with a heritability of 0.3 represent production traits (i.e., daily gain, feed intake, feed efficiency, back fat thickness, and muscle depth) (Bergsma *et al*., 2013; Godinho *et al*., 2018; Ogawa *et al*., 2022). Insight into the selection history of traits known to be under selection can be a valuable addition to current practices, where the impact of selection is generally only assessed based on the genetic trend, which shows how the average estimated breeding values change across generations. Therefore, selection history information can aid in the evaluation of the effectiveness of the applied selection index and whether or not selection weights should be adjusted.

Additionally, we were interested in the potential to investigate the selection history of new traits, which might be interesting to add to the breeding goal. This is because the focus of livestock breeding goals is continuously shifting towards more balanced breeding. Hence, selection is not only on production and reproduction traits, but novel traits related to robustness, resilience, welfare, environmental footprint, disease resistance, and social behaviour have been or are currently being included in the breeding goal (Kanis *et al*., 2005; Merks *et al*., 2012; Harlizius *et al*., 2020). These new traits tend to have lower heritabilities because they are more complex, influenced by many genes and by the environment (Rydhmer and Lundeheim, 2008; Turner *et al*., 2024). If we want to determine whether there has been indirect selection on these traits by selection on the breeding goal, the selection accuracy for these traits can be expected to be weak. Using lower heritability values and phenotypic selection in our simulations, therefore, provides a realistic and more challenging scenario for selection history models. Information on the selection history of those traits can indicate whether there are strong genetic correlations with other breeding goal traits, which is important to know before the trait is included in the breeding goal.

### Relation between the effect size and MAF under selection

In populations under natural selection, negative (purifying) selection acts against mutations with large deleterious effects and prevents them from becoming frequent in a population (Pritchard, 2001; Eyre-Walker, 2010). These deleterious mutations are kept at low frequency in the population by the principles of mutation-selection balance, resulting in the negative relationship between MAF and effect size, as is often observed in human populations (Speed *et al*., 2017, 2020; Timpson *et al*., 2018; Zeng *et al*., 2018, 2021). In contrast, directional (positive) selection acts to increase beneficial alleles in the population, potentially leading them to fixation (Walsh and Lynch, 2018). This would initially result in a positive relationship between the MAF and effect size, where the positive effect alleles are more frequent at intermediate frequencies. Therefore, a negative relation between MAF and effect size in populations under natural selection indicates purifying selection, whereas a positive relation indicates directional selection.

In populations under artificial selection, the estimated *s-*values should be interpreted differently than in populations under natural selection. This is because populations under artificial selection often have a smaller effective population size. In those populations, directional selection can result in a rapid increase in frequency of beneficial alleles until they reach fixation (Rowan *et al*., 2021), because the selection pressure is high on large-effect loci, resulting in a rapid increase in the allele frequency of these loci. Therefore, alleles with a large effect only remain shortly at intermediate allele frequencies and those large-effect loci have on average a low MAF. The selection pressure on loci with small effects is relatively low, resulting in slower fixation and therefore they have, on average, a higher MAF. Hence, in breeding populations under directional selection, a negative relationship between MAF and effect size is expected. This negative relationship has been observed for important breeding goal traits in tilapia, dairy cattle, maize, and pigs (Joshi *et al*., 2021; Reynolds *et al*., 2022; Palali Delen *et al*., 2023; Neshat *et al*., 2023; Zhang *et al*., 2025). This is in agreement with the results of our study, where we found a negative relationship between the effects and the MAF of the causal loci after 30 generations of directional selection (Figure 10), which was correctly reflected in the estimated *s-*values.

Overall, we have seen that the optimal *s*-values became more negative when the number of generations of ongoing selection increased for a trait under directional selection (Figure 8). However, when we look at the ΔAIC_s_ scores over these generations for a heritability of 0.3, ΔAIC_s_ scores >10 compared to zero occurred in 0%, 2%, 6%, 6%, 12%, 4%, 0% of the replicates for generation 1, 5, 10, 15, 20, 25, and 30, respectively (Appendix III). So, at first there was an increase in power up to generation 20, where selection was most often detected, after which there was a decrease in power, resulting in a decline in the detection of selection. As no new mutations were simulated during selection, more alleles reached fixation as the number of selected generations increased. This probably decreased the power to detect selection for a heritability of 0.3. For the lower heritabilities, the fixation of alleles occurred at a slower rate caused by the lower accuracy of selection, and therefore, we don’t see this pattern there.

### Bias of *s*-value estimations

For random selection, the MAF and effect size were expected to be independent from each other, resulting in an expected optimal *s-*value of zero. Indeed, the regression coefficient for the MAF and the effect size of the causal loci after 30 generations of selection after the burn-in was close to zero (0.02). However, the optimal *s-*values were, on average, estimated with SNPs to be around -0.2 to -0.3 for random selection. First, we investigated the optimal *s*-value using the causal loci as input, which resulted in an *s*-value of 0.1 (Figure 9). This indicates that the expectation for an *s*-value of approximately zero is valid based on the causal loci only. Second, we investigated whether the negatively estimated *s*-values were a result of ascertainment bias, as the SNPs were sampled to have a uniform allele frequency distribution, while the causal loci had a U-shaped allele frequency distribution. This could result in not being able to explain the causal loci with low MAF accurately by the SNP, as is a common bias for SNP arrays (Nielsen, 2004; Clark *et al*., 2005; Geibel *et al*., 2021). However, even with SNPs from a U-shaped allele frequency distribution, the optimal *s*-value was -0.3 for random selection with a heritability of 0.3 (Figure 9), so the ascertainment bias was not the reason for finding an estimated *s*-value below zero. Overall, the ΔAIC scores for the optimal *s*-values for random selection reveal that for a maximum of 2% of the replicates, a value of ΔAIC_s_>10 is found when comparing the optimum *s*-value to a value of zero (Appendix II). The *s*-values are thus estimated as slightly more negative based on SNPs than expected based on causal loci, but only rarely deviated significantly from zero.

For directional selection, the *s*-values were estimated to be -1 for both SNP arrays (uniform and U-shaped allele frequency distribution), whereas the *s*-value based on the causal loci was estimated to be -0.7. Similar to what was seen for random selection, the estimated *s-*values based on SNPs seem to be slightly more negative than the estimated *s*-values based on causal loci. This underestimation of the *s*-value could be due to linkage disequilibrium (LD) between the causal loci and the SNPs, which is often incomplete and not all causal loci can be completely explained by the SNP. Moreover, not all SNPs might be linked to a causal locus under selection. It is good to note that we have applied phenotypic selection, so the SNPs were not used for the selection. Therefore, it is expected that the causal loci reflect the true relationship between effect size and minor allele frequency in the population. So, the *s*-values estimated from SNPs are more negative than from causal loci, thereby overestimating the actual selection signal at the causal variants.

### Comparing *s*–value estimation and Ĝ

We have applied two models to investigate selection history to evaluate their ability to detect selection in an animal breeding population under ongoing directional selection. The *s-*value estimation has been mostly applied to human populations to identify selection (Speed *et al*., 2017, 2020; Zeng *et al*., 2018, 2021), and the interpretation of the *s-*values is slightly different for artificial selection as explained before. Ĝ, on the other hand, is a model based on principles of genomic selection and has been applied to breeding populations (Beissinger *et al*., 2018).

When applying Ĝ in a population under directional selection, we have seen that for a selection interval of 10 generations, correct detection of selection occurred in 30%, 56%, and 98% of the replicates for a trait with a heritability of 0.05, 0.1, and 0.3, respectively. For each selection scenario this percentage increased when the selection interval became longer. The detection of selection for a heritability of 0.3 was nearly perfect for a selection interval of at least 5 generations. For a heritability of 0.1, selection was detected in at most 70% of the replicates, which occurred for a selection interval of 15 generations. For a heritability of 0.05 selection was identified at most for 38% of the replicates for a selection interval of 20 generations. In general, increasing the selection interval beyond 10 generations did not result in a large increase in detection of selection. This could be explained by the fact that marker effects are not constant over time. Ongoing selection changes allele frequencies, which in turn can alter linkage disequilibrium patterns and the additive genetic variance. As a result, the estimated marker effects may change over generations (Wientjes *et al*., 2022). Extending the selection interval too far may, therefore, limit further improvement in detection, as ongoing allele frequency changes alter the underlying genetic architecture. From this, we can conclude that Ĝ detects selection reasonably well for low heritability traits, where only detection for a trait with a heritability of 0.05 was limited, although we do see small improvements when the selection interval is increased.

Another way to improve the detection of selection is potentially to increase the number of genotyped animals in the start and end generations, as shown by Mahmoud *et al*. (2023). Implementing this in practice would require that allele frequency data from 10 to 20 generations back in time would have to be available for these animals to potentially increase the detection of selection with Ĝ for a heritability of 0.05. Alternatively, initial allele frequencies could be inferred from pedigree data (Gengler *et al*., 2007), especially if the main parents of the investigated generations are genotyped. However, it has been described that this likely affects the detection of selection by reducing the power of the model (Beissinger *et al*., 2018).

Different from Ĝ, the estimation of *s*-values uses only data from the most current generations. Estimating *s*-values as an indication of selection resulted in the correct identification of selection of 4%, 46%, and 100% for heritabilities of 0.05, 0.1, and 0.3, respectively, when the *s*-value was estimated using the last five generations of selection (Table 3). Increasing the number of generations, and thereby the number of genotyped animals, to which the model is applied, increased the detection of selection. Our study has shown that using the last five generations of the breeding population greatly increased the detection of selection compared to the last generation only, also for lower heritabilities.

The additive marker effects that are used to determine the relationship between the MAF and the effects can be more accurately estimated as the sample size increases (Daetwyler *et al*., 2008). So, enlarging the dataset further with more genotyped and phenotyped animals of one or more generations is expected to increase the correct identification of selection for the lower heritabilities more. However, it is good to realize that those marker effects can fluctuate over time as allele frequencies change under selection, thereby altering the LD patterns and the apparent size of marker effects (Goddard, 2009). So, increasing the number of animals in the last generations can result in a more accurate estimation of the marker effects in the last generation, but increasing the number of animals across generations to estimate markers effects can help to estimate marker effects across generations that might be more related to the applied selection. Therefore, it is difficult to determine based on our results whether increasing the number of animals in the last generations or across generations contributes more strongly to improved detection. This should be considered when applying this model to empirical data.

In summary, considering both the strengths and weaknesses of each model, we can say that Ĝ is an intuitive model for breeders and the interpretation of positive or negative selection is straight forward, but the ability to reliably detect selection may be limited by the availability of the required data. Ĝ has shown to very accurately detect selection for traits with a moderate heritability (0.3). For lower heritabilities (0.05 and 0.1) the detection of selection is a bit lower but still substantial. Increasing the selection interval improved the detection of selection, but increasing the number of genotyped animals could possibly improve the results further. However, it is good to realize that the allele frequency data of animals from 10 or more generations ago might not always be available, or not in the numbers required to detect selection. In contrast to Ĝ, the *s*-value method has no clear significance value and therefore the identification of selection is a bit more arbitrary. Furthermore, no distinction can be made between positive and negative selection, as both result in a negative value for *s* in populations under strong artificial selection. All in all, we have seen that both models are capable of detecting selection in animal breeding populations, but there are different requirements for the dataset to reliably detect selection.

## Conclusion

In conclusion, both Ĝ and the *s*-value estimation are able to detect selection in a simulated pig breeding population for traits with low to medium heritabilities. In general, higher heritabilities and a larger sample size (for *s*-value estimation) or a larger selection interval (for Ĝ) result in better detection of selection. It depends on the available genotype- and phenotype information and the trait of interest, which model would be best to apply. In general, Ĝ would be the preferred model to apply when allele frequency data from at least 10 generations back is available for a population, while the *s*-value estimation is the preferred model when only the latest generations of a population have been genotyped.

## Author Contributions

MPLC and YCJW obtained funding for this study. ACMJ, MPLC, and YCJW participated in the design of the study. ACMJ performed the analyses and drafted the manuscript. ACMJ, MPLC, and YCWJ were involved in the interpretation of the results. All authors read and approved the final manuscript.

## Data Availability Statement

All scripts used for simulations and analysis are available at Figshare, in the folder scripts_selection_history_models.

## Appendix I. Δ𝑨𝑰𝑪_𝒔_ scores *s-*value estimation directional selection

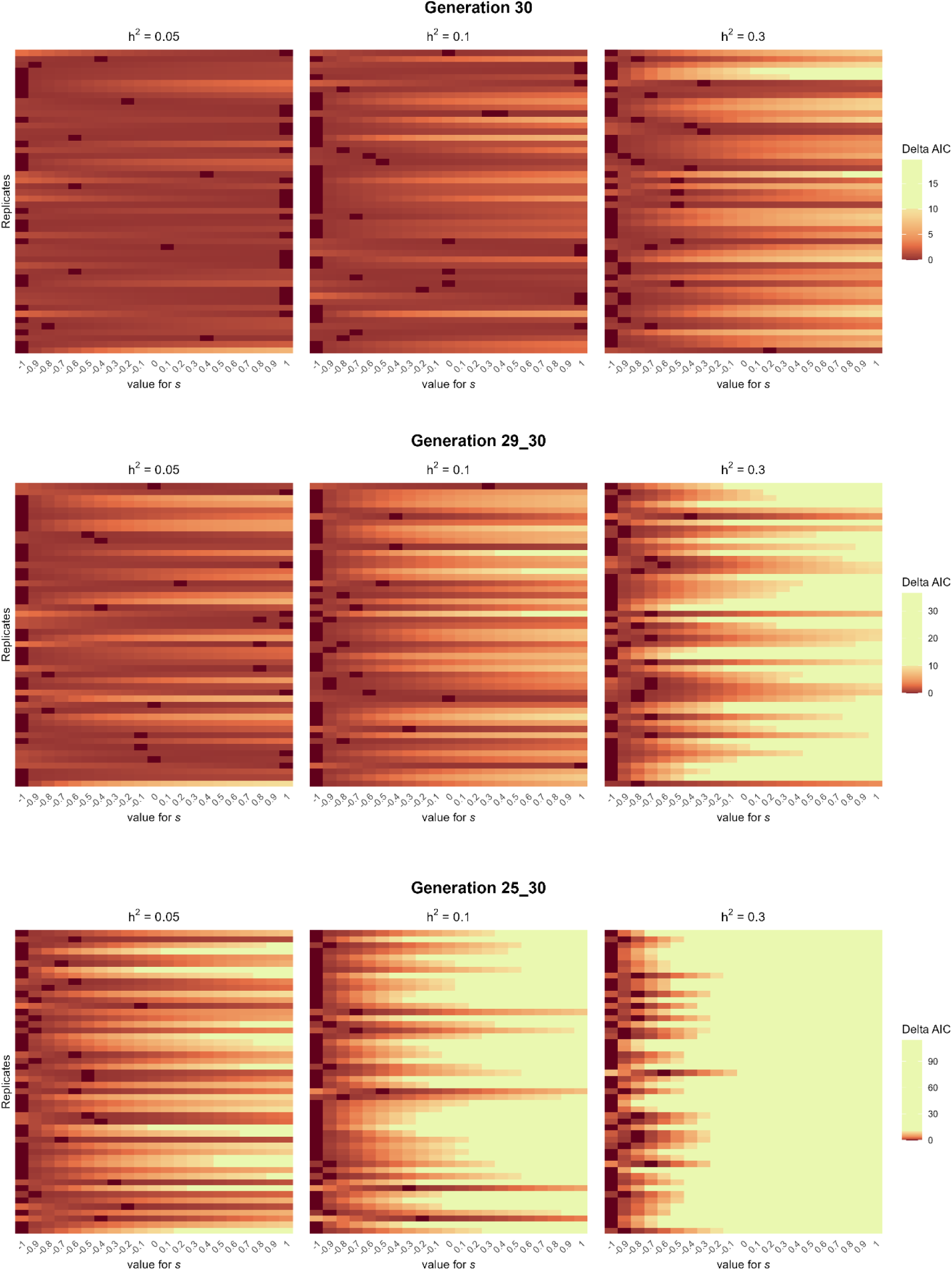

## Appendix II. Δ𝑨𝑰𝑪_𝒔_ scores *s-*value estimation random selection

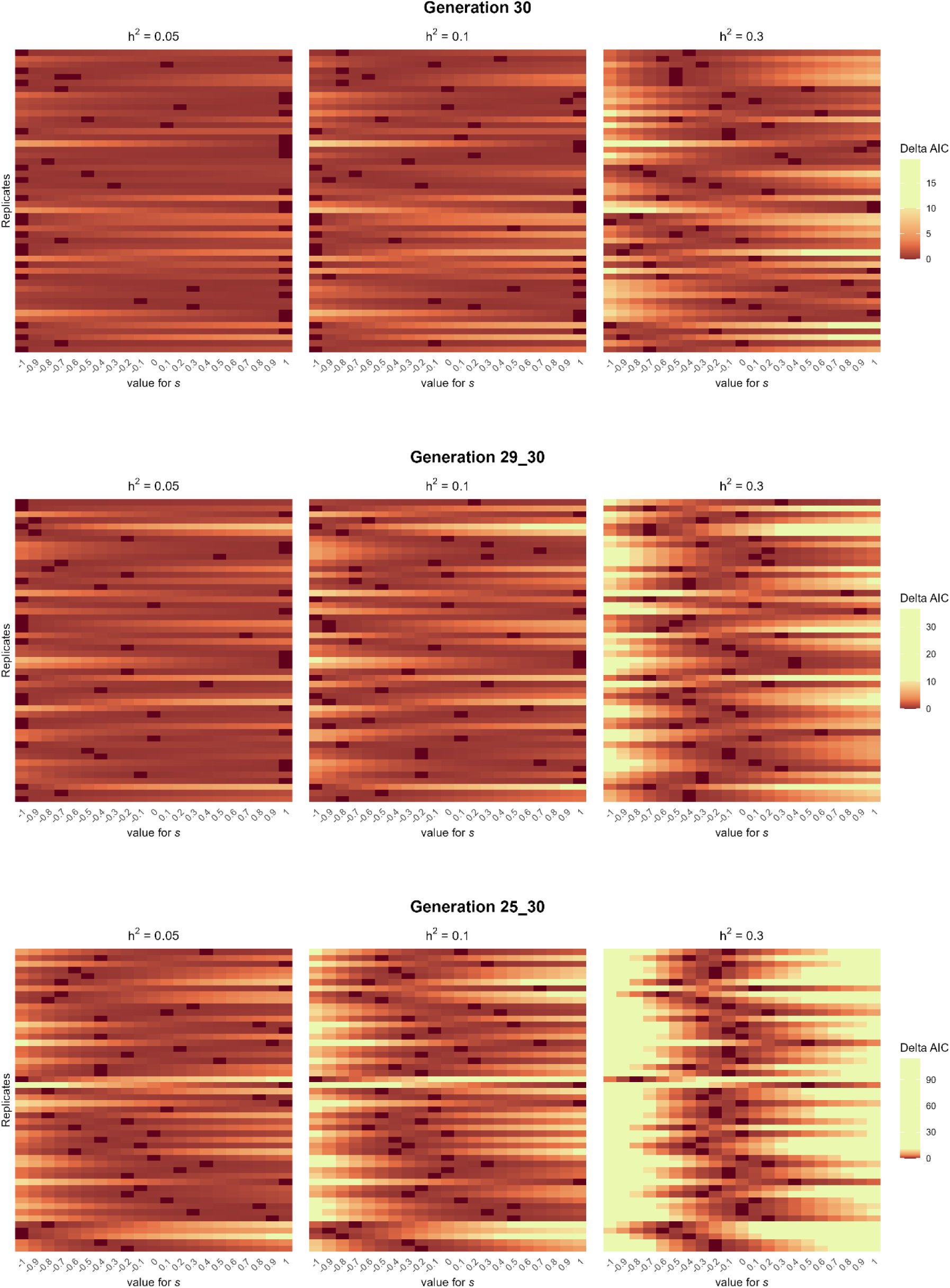

## Appendix III. Δ𝑨𝑰𝑪_𝒔_ scores *s-*value estimation directional selection extra generations

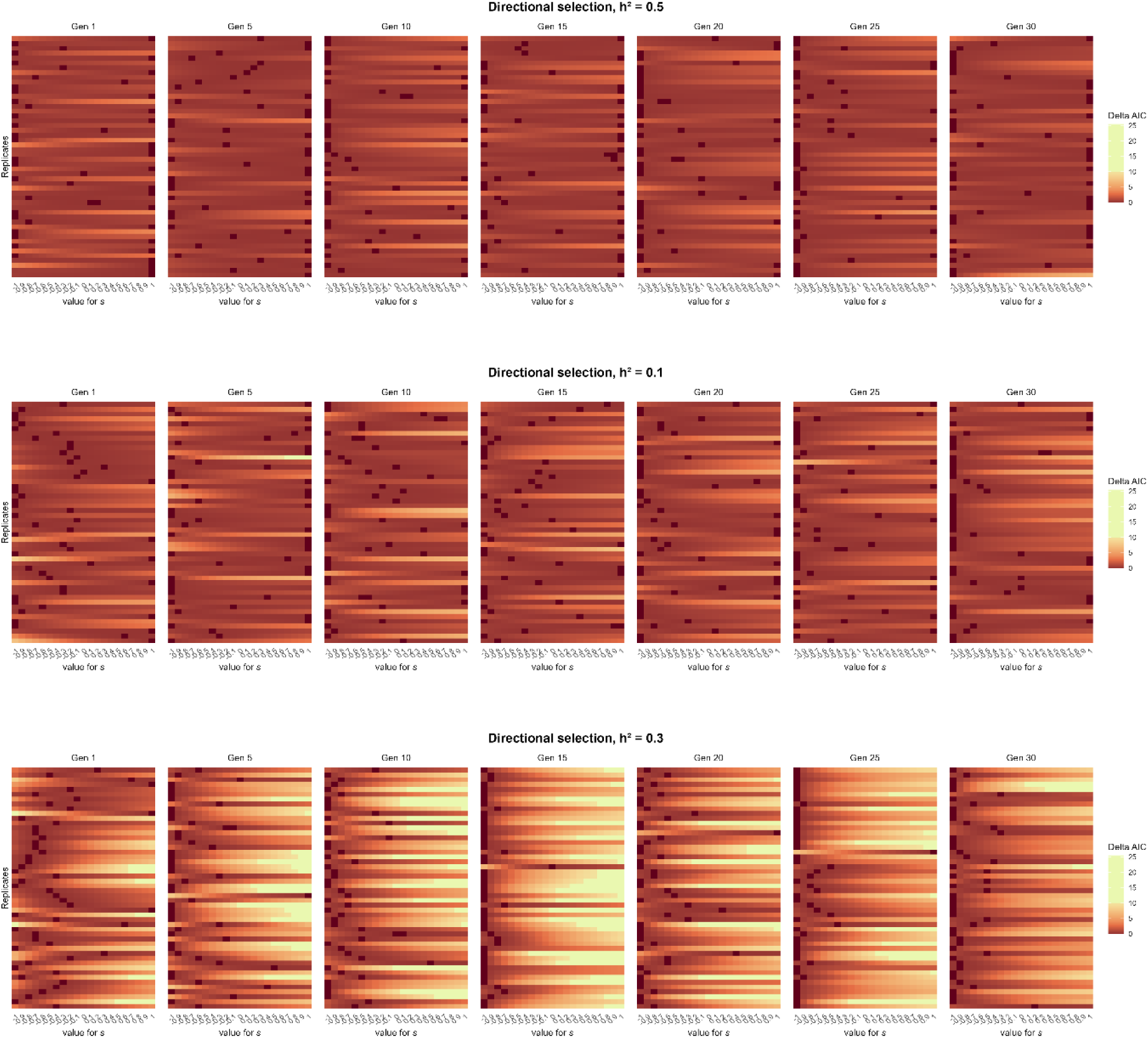

